# Single Transcript Level Atlas of Oxytocin and the Oxytocin Receptor in the Mouse Brain

**DOI:** 10.1101/2024.02.15.580498

**Authors:** Vitaly Ryu, Anisa Gumerova, Georgii Pevnev, Funda Korkmaz, Hasni Kannangara, Liam Cullen, Ronit Witztum, Steven Sims, Tal Frolinger, Ofer Moldavski, Orly Barak, Jay J. Cao, Daria Lizneva, Ki A. Goosens, Tony Yuen, Mone Zaidi

## Abstract

Oxytocin (OXT), a primitive nonapeptide known to regulate reproduction and social behaviors, is synthesized primarily in the hypothalamus and is secreted *via* hypophyseal-portal system of the posterior pituitary gland. Given that pituitary hormones, traditionally thought of as regulators of single targets, display an array of central and peripheral actions, OXT also directly affects bone and body composition. Its effects on bone remodeling are physiologically relevant, as elevated OXT levels during pregnancy and lactation could cause calcium mobilization from the maternal skeleton for intergenerational calcium transfer towards fetal bone growth. There is an equally large body of evidence that has established the presence of OXT receptors (OXTRs) in the brain through which central functions, such as social bonding, and peripheral functions, such as the regulation of body composition, can be exerted. To purposefully address the effects of OXT on the brain, we used RNAscope to map OXT and OXTR expression, at the single transcript level, in the whole mouse brain. Identification of brain nuclei with the highest OXT and OXTR transcript density will shed further light on functional OXT nodes that could be further interrogated experimentally to define new physiologic circuitry.

## INTRODUCTION

Oxytocin (OXT), a neuropeptide synthesized primarily by magnocellular neurons within the paraventricular (PVH) and supraoptic nuclei (SON) of the hypothalamus (Landgraf & Neumann, 2004; Sofroniew, 1983; Swanson & Sawchenko, 1983), has been broadly implicated in the control of parturition, lactation, appetite, emotions, stress responses, and social behavior. The distribution of OXT receptors (OXTRs) across the brain in different species provides a proxy for the distribution of OXT binding, thus providing evidence for OXT nodes in the brain of physiologic relevance. It has been reported that OXTRs are heterologously expressed in many brain sites, including the central nucleus of the amygdala and the ventromedial hypothalamic nucleus (VMH) (Bale & Dorsa, 1995a, b). Furthermore, *Oxtr* mRNA has been detected in the hypothalamus, olfactory bulb, ventral pallidum, and the dorsal vagal nucleus (Adan *et al*, 1995; Yoshimura *et al*, 1993).

OXT mediates a variety of peripheral and central functions. While the peripheral actions comprise milk ejections, uterine contractions, and prolactin production, the central actions of OXT are mostly related to female reproduction, including sexual receptivity (Caldwell *et al*, 1986), pair ponding (Insel, 1992) and maternal behavior (Fahrbach *et al*, 1984; Insel, 1990; Pedersen *et al*, 1982). Central functions of OXT also include modulation of cardiac vagal input (Bohus *et al*, 1996), memory consolidation (Dyball & Paterson, 1983) and social/affiliative behavior (Insel, 1992; van Wimersma Greidanus & Maigret, 1996). Axons and dendrites of OXT neurons are localized in close proximity to the third ventricle and even in between tanycytes and ependymal cells facing the cerebrospinal fluid (Landgraf & Neumann, 2004). Notably, magnocellular OXT neurons send extended dendritic trees, forming the basis for the somato-dendritic release of OXT within the PVH and SON (Ludwig & Leng, 2006; Neumann *et al*, 1993; Neumann, 2007; Pow & Morris, 1989). Such release is likely to facilitate autocrine and/or paracrine regulation of OXT neurons towards physiologic demands, such as lactation (Moos & Richard, 1989; Neumann *et al*, 1994) and child birth (Neumann *et al*, 1996). To exert neuronal effects, locally released OXT binds to local OXTRs, which are expressed within or are juxtaposed to the target region, for example, on synapses, as well as on axons and glial processes (Mitre *et al*, 2016). Alternatively, OXT could putatively diffuse over longer distances to bind to adjacent OXTRs (Landgraf & Neumann, 2004; Ludwig & Leng, 2006; Mitre *et al*., 2016). Given OXT exerts its multiple behavioral effects through its action on several regions of the forebrain and mesolimbic brain, the question whether other extrahypothalamic projections of OXT neurons may also have a role garners significant importance.

It is also becoming increasingly clear that both anterior and posterior pituitary hormones, traditionally thought of as regulators of single physiological processes, affect multiple bodily systems, either directly or *via* actions on brain receptors (Abe *et al*, 2003; Zaidi *et al*, 2018). Non-traditional actions of OXT include its ability to affect the skeleton, wherein it stimulates bone formation by osteoblasts and modulates the function of bone-resorbing osteoclasts (Sun *et al*, 2019). We have shown that OXT and vasopressin have opposing skeletal actions-effects that may relate to the pathogenesis of bone loss in pregnancy and lactation, and in chronic hyponatremia, respectively (Sun *et al*., 2019; Sun *et al*, 2016; Tamma *et al*, 2009; Tamma *et al*, 2013). We have also noted that regions known to send sympathetic nervous system (SNS) outflow to both bone and adipose tissues express the *Oxtr* (Ryu *et al*, 2015; Ryu *et al*, 2017). This lends to the possibility that certain actions of OXT on peripheral tissues, such as on body composition and bone, may also be mediated centrally.

Despite a *corpus* of evidence for the expression of OXT and OXTRs in various brain regions, and their function in regulating central and peripheral actions, such as social behavior and satiety (Bale *et al*, 2001; Sun *et al*., 2019), there remains the need for a detailed, gender-specific mapping of the anatomical geography of the OXT system in the brain. Here, we use RNAscope-a cutting-edge technology that detects single RNA transcripts-to create a comprehensive atlas of the OXT and OXTR in the mouse brain. We believe that this compendium of OXT and its receptor in concrete brain sites at a single transcript level should provide a resource for investigators to study both peripheral and central effects of interrogating OXTRs site-specifically in health and disease. Our identification of brain nuclei with the highest OXT and OXTR transcript density will thus deepen our future understanding of the functional engagement of the central OXT-containing neuronal nodes within a large-scale functional network.

## RESULTS

Mapping autoradiographic studies suggest that the distribution of OXTRs in the brain varies greatly among different rodent species (Dubois-Dauphin *et al*, 1992; Elands *et al*, 1988; Insel *et al*, 1997; Tribollet *et al*, 1992). Besides mapping the full anatomical distribution of *Oxt* and *Oxtr* by RNAscope, the present study also assessed sex differences in *Oxt* distribution. Allowing the detection of single transcripts, RNAscope uses ∼20 pairs of transcript-specific double Z-probes to hybridize 10-µm-thick whole brain sections. Preamplifiers first hybridize to the ∼28-bp binding site formed by each double Z-probe; amplifiers then bind to the multiple binding sites on each preamplifier; and finally, labeled probes containing a fluorescent molecule bind to multiple sites of each amplifier.

RNAscope data were quantified on sections from coded mice. Each section was viewed and analyzed using CaseViewer 2.4 (3DHISTECH, Budapest, Hungary) and QuPath v.0.2.3 (University of Edinburgh, UK). The *Atlas for the Mouse Brain in Stereotaxic Coordinates* (Paxinos & Franklin, 2007) was used to identify every nucleus, sub-nucleus or region, which was followed by manual counting of *Oxt* and *Oxtr* transcripts in every tenth section using a tag feature. Receptor density was calculated by dividing the transcript number by the area (µm^2^, ImageJ) in every nucleus, sub- nucleus or region. Photomicrographs were prepared using Photoshop CS5 (Adobe Systems) only to adjust brightness, contrast and sharpness, and to remove artifacts (i.e., obscuring bubbles).

We report the expression of the *Oxtr* in 359 mouse brain nuclei, sub-nuclei and regions. Probe specificity was established by a positive signal in the epididymis with an absent signal in the liver (negative control) (Fig. 1A). Notably, *Oxtr* transcripts were detected bilaterally, with no apparent ipsilateral domination. Transcripts were highest in the ventricular regions followed, in descending order, by the hypothalamus, olfactory bulb, hippocampus, cerebral cortex, medulla, midbrain and pons, forebrain, thalamus and cerebellum (Fig. 1B). Using the RNAscope dataset we further calculated *Oxtr* density in various brain nuclei, sub-nuclei and regions. High *Oxtr* transcript counts and densities, respectively, were also noted in several nuclei, sub-nuclei and regions as follows (Fig. 1C): ventricular regions-ependymal region for both; hypothalamus- AHiPM for both; olfactory bulb-vn and GrO; hippocampus-SLu and Py; cerebral cortex-CI and Pir; medulla-10N and Sp5I; midbrain and pons-IPF and CIC; forebrain-aci and CPu; thalamus-PV and PVA and cerebellum-7Cb and Sim [see Appendix for nomenclature and Supplementary Fig. 1 for transcript count and representative photomicrographs].

**Figure 1:**
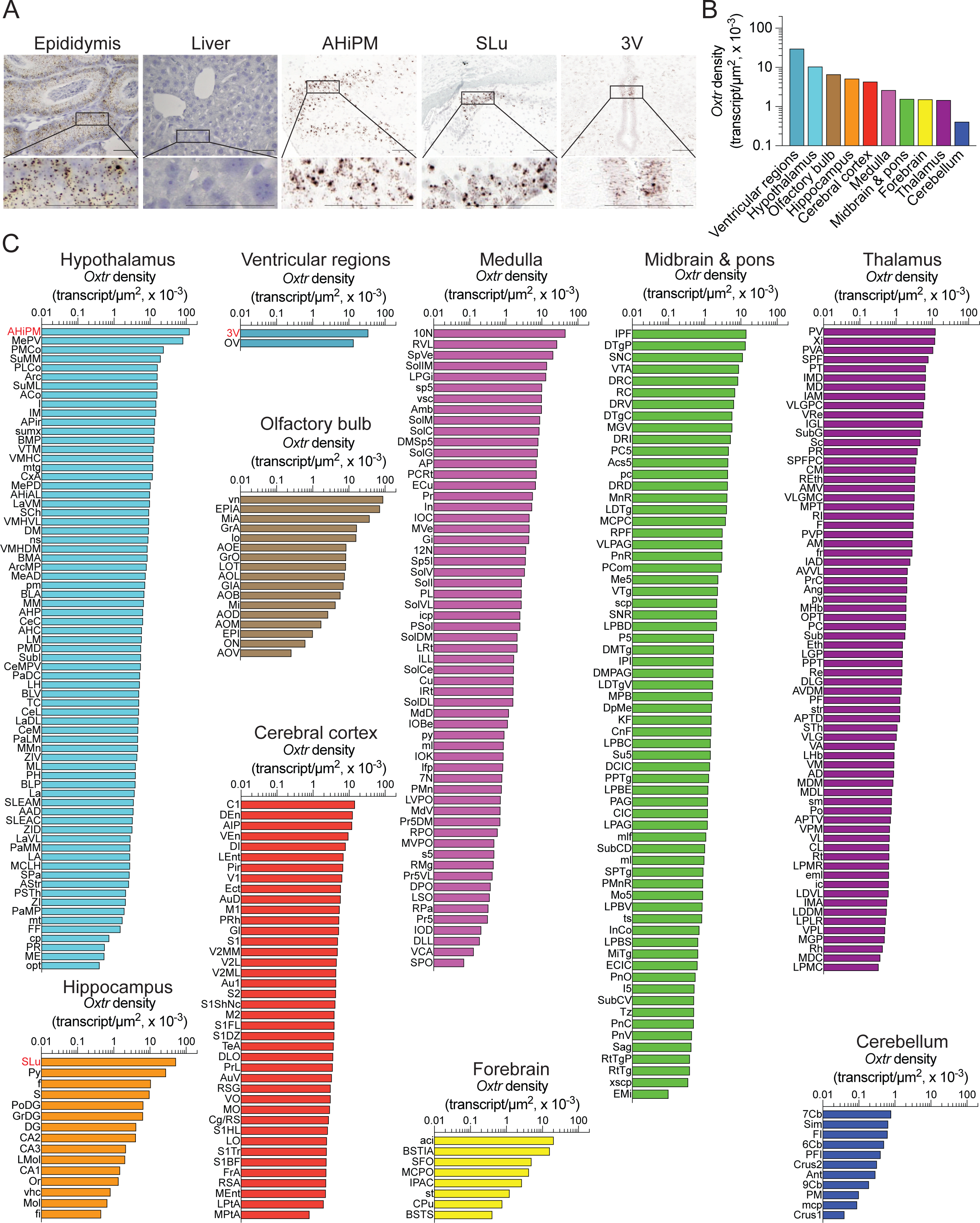
*Oxtr* expression in the mouse brain. (**A**) RNAscope revealed *Oxtr*-positive transcripts in the epididymis, but not in the liver (positive and negative controls, respectively). Also shown are representative micrographs in the posteromedial part of the amygdalohippocampal area (AHiPM) of the hypothalamus, stratum lucidum (SLu) of the hippocampus, and the third ventricle (3V) of the ventricular regions. Scale bar: 100 µm. (**B**) *Oxtr* transcript density in the brain regions detected by RNAscope. (**C**) *Oxtr* transcript density in nuclei, sub-nuclei and regions of the ventricular system, hypothalamus, olfactory bulb, hippocampus, cerebral cortex, medulla, midbrain and pons, forebrain, thalamus, and cerebellum.

RNAscope also revealed *Oxt* expression in the hypothalamus and forebrain of both male and female mice (Figs. 2A, 2B). High *Oxt* counts were detected in several nuclei, sub-nuclei and regions of females and males, respectively, as follows (Fig. 2C): hypothalamus-PaMP and PaMM and forebrain-MPA and LPO. Overall, the numbers of *Oxt*-expressing cells were markedly greater in female compared to the male mice. That is, we found 22 hypothalamic and forebrain regions with 486 *Oxt*-positive cells in the female mouse. In contrast, there were 15 hypothalamic and forebrain regions containing 308 *Oxt*-positive cells in the male brain. Breaking this down, in the hypothalamus, the number of *Oxt*-positive cells was 372 in the female compared with 228 *Oxt*-positive cells in the male. The number of *Oxt*-positive cells was 114 in the female forebrain compared with 80 *Oxt*-positive cells in the male forebrain.

**Figure 2:**
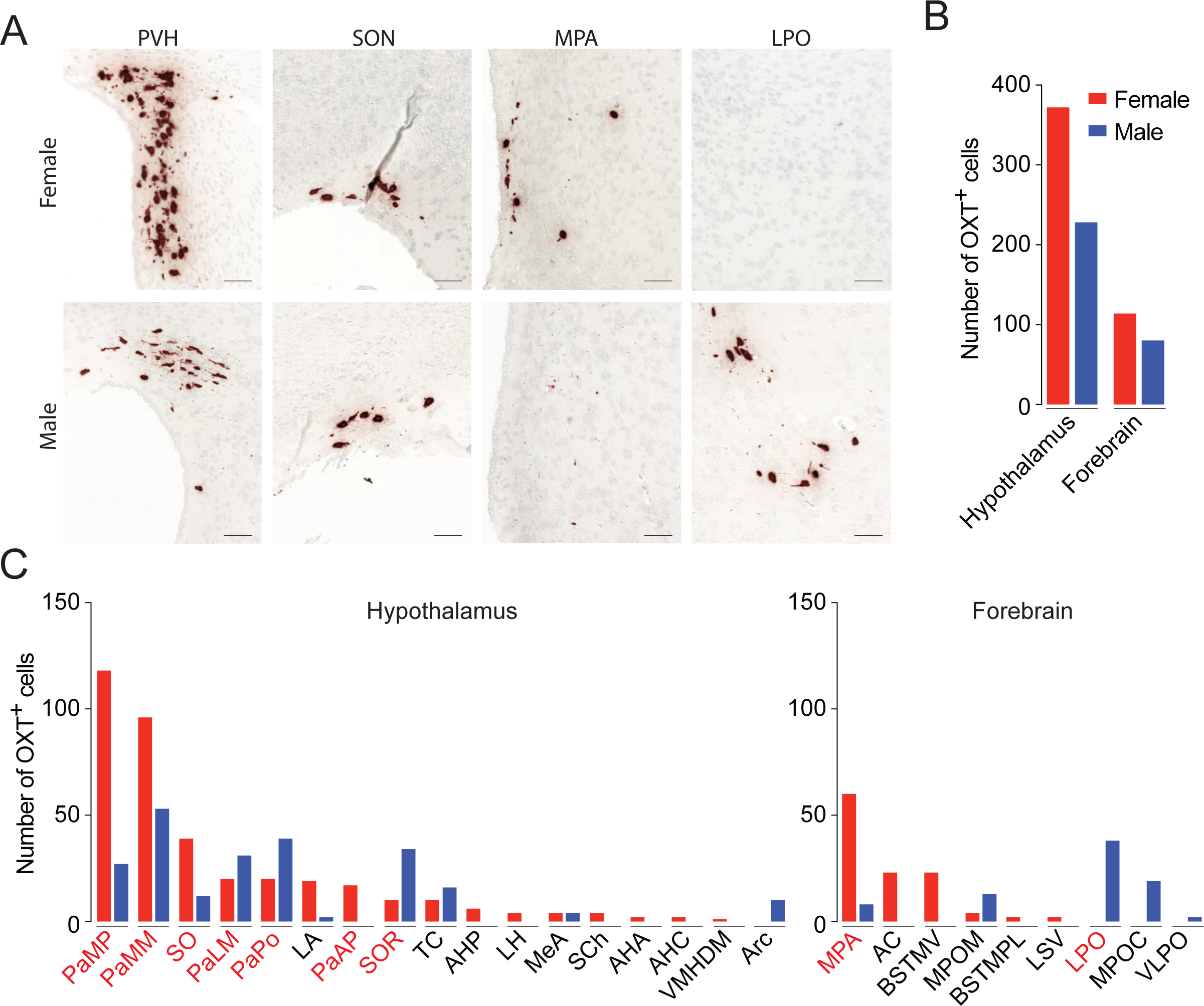
Gender differences of *Oxt* expression in the mouse brain. **(A)** The neurohypophysial hormone OXT is synthesized by magnocellular neurons primarily located in the PVH and SON hypothalamic nuclei. The magnocellular neurons send extended axonal projections into the neurohypophysis where OXT is released in to the circulation in response to physiological demands. Therefore, PVH and SON served as positive controls for OXT expression in the brain. MPA: medial preoptic area; LPO: lateral preoptic area of the forebrain. Scale bar: 50 µm. (**B**) Gender differences in the numbers of *Oxt-*expressing neurons in nuclei, sub-nuclei and regions of the hypothalamus and forebrain detected by RNAscope. (**C**) Total numbers of *Oxt-* expressing neurons in nuclei, sub-nuclei and regions of the hypothalamus and forebrain of male and female mice.

*Oxtr* expression was also mapped in regions and sub-regions within the hypothalamus (Fig. 1 and Supplementary Fig. 1). Certain of these hypothalamic subregions, such as the lateral hypothalamus (LH) and dorsomedial hypothalamus (DM), send SNS outflow to both bone and fat tissue (Ryu *et al*., 2015; Ryu *et al*., 2017). Additionally, RNAscope also showed *Oxtr* expression in both anterior and posterior pituitary lobes (Fig. 3A), with *Oxtr* transcript density that was markedly higher in the female compared with male mice (Fig. 3B).

**Figure 3:**
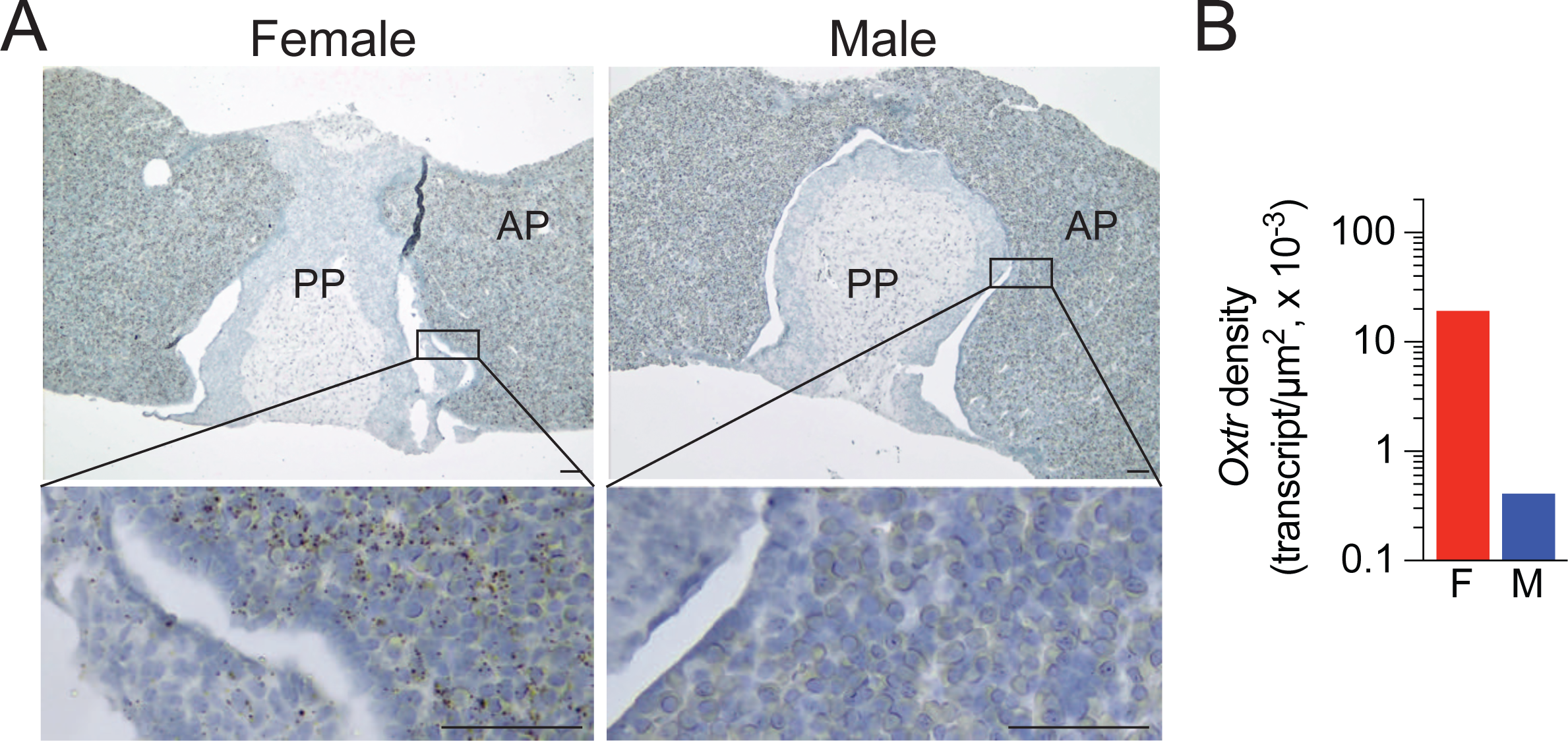
Sex differences in *Oxtr* expression in the pituitary gland. **(A)** Representative photomicrographs showing sex differences in *Oxtr* expression in anterior (AP) and posterior (PP) lobes of the pituitary gland detected by RNAscope. Scale bar: 25 µm. (**B**) Quantification of *Oxtr* transcript density in the pituitary gland of female and male mice.

We also found that six hypothalamic nuclei, sub-nuclei and regions in male mice displayed overlapping *Oxt* and *Oxtr* transcripts. *Oxtr*/*Oxt* ratios within the same brain site were as follows: 0.70 in the medial parvicellular part of the paraventricular hypothalamic nucleus (PaMP); 1.51 in the medial magnocellular part of the paraventricular hypothalamic nucleus (PaMM); 6.39 in the lateral magnocellular part of the paraventricular hypothalamic nucleus (PaLM); 26.80 in the arcuate nucleus (Arc); 109.50 in the medial amygdaloid nucleus (MeA); 151.13 in the tuber cinereum area (TC), and 222.00 in the lateroanterior hypothalamic nucleus (LA).

## DISCUSSION

Here, we supplement and integrate previous information on OXT and OXTR expression in the murine brain and report, for the first time, abundant OXTR expression in 359 brain nuclei, sub-nuclei and regions, as well as, importantly, gender-specific *Oxt* expression. This report is thus the most comprehensive atlas of brain *Oxt* and *Oxtr* expression at the single transcript level. Expression of both *Oxt* and *Oxtr*, particularly in overlapping hypothalamic sub-nuclei, nuclei, and regions, points to functionally active neuronal nodes within a large-scale OXT-OXTR network in the brain.

It has been reported that cell bodies and dendrites of OXT-producing neurons within the PVH and SON release OXT and AVP within the magnocellular nuclei, where we find the highest *Oxtr* expression-this suggests an additional, possibly paracrine action of OXT (Ludwig & Leng, 2006; Neumann *et al*., 1993; Neumann, 2007; Pow & Morris, 1989). Indeed, locally released OXT is involved in pre-and post-synaptic modulation of the electrical activity (Bourque *et al*, 1993; Kombian *et al*, 1997; Shibuya *et al*, 2000). Similar to magnocellular neurons of the PVH, we find that several *Oxt-* producing neuronal populations also overlap with *Oxtr* expression in other hypothalamic Sites—hereby termed “ *Oxtr/Oxt* nodes” —these include Arc, MeA, TC and LA.

Of note is that, in addition to the nucleus of the solitary tract (NTS), the *Oxtr*/*Oxt* node in the Arc (and, possibly, Arc-bordering LA) rapidly induces satiety (Fenselau *et al*, 2017) and suppresses excessive food intake to control ingestive behavior (Inada *et al*, 2022). The *Oxtr*/*Oxt* node in the MeA likely explains the paracrine regulation by OXT of male preference for for females and their scents (Yao *et al*, 2017). Thus, the ablation of *Oxtr* in aromatase-expressing neurons of the MeA fully recapitulates the elimination of female preference in males, suggesting that this node is both necessary and sufficient for social recognition (Ferguson *et al*, 2001). Lastly, the TC is a sheet of grey matter that forms a median eminence (ME) around the base of the pituitary stalk or infundibulum: therefore, the *Oxtr*/*Oxt* node in the TC (and tanycyte) could be important for mediating bidirectional brain-periphery crosstalk by modulating the blood-hypothalamus brain barrier.

Surprisingly, the highest *Oxtr* transcript density was noted in the tanycyte-rich ependymal layer and, not surprisingly, in the hypothalamus (Landgraf & Neumann, 2004; Sofroniew, 1983; Swanson & Sawchenko, 1983). In the hypothalamus, the highest density was found in the posteriomedial part of the amygdalohippocampal area (AHiPM). It has been reported that ∼40% of *Oxtr-*positive neurons of the amygdalohippocampal area (AHi) project to the medial preoptic area (MPOA) (Sato *et al*, 2020). Activation of these neurons, comprising excitatory projections to the MPOA, enhances exclusively an aggressive, but not parental behavior, towards pups (Sato *et al*., 2020).

The olfactory bulb and hippocampus also displayed abundant *Oxtr* transcripts, with the highest density in the vomeronasal nerve (vn) and the ventral stratum lucidum (SLu), respectively. It has been reported that *Oxtrs* are expressed in the vomeronasal organ, an olfactory sensory structure involved in the detection of non-volatile chemosignals. OXT injection in mice has been shown to reduce pup-induced vomeronasal activity and aggressive behavior (Nakahara *et al*, 2020). Given vomeronasal activity declines as males grow up from a pup-aggressive state to a non-aggressive parental state, high *Oxtr* expression in the vn might indicate a functional switch from pup-aggressive behavior towards strengthening social and sexual behaviors during adolescence and adulthood. As with the hippocampal SLu, *Oxtrs* are found in both excitatory and inhibitory neurons within the CA2 and CA3 subregions of the hippocampus, suggesting that OXTRs in the SLu may have a role in local circuits relating to stress, emotional and affective behaviors.

In the cortex, we found that C1 adrenaline cells displayed the highest *Oxtr* transcript density. These cells project to the SON, and upon activation by the OXTR release OXT from the SON-a putative paracrine loop. Moreover, OXTRs mediate cardiac sympathetic stimulation through direct PVH projections to the intermediolateral column of the spinal cord (Japundzic-Zigon, 2013). Such reciprocal communications are supported by the studies inferring that affiliative social interactions increase OXT activity, which is followed by an anti-stress response, thus promoting bonding, relaxation and growth, while reducing cardiovascular and neuroendocrine stress (Grippo *et al*, 2009; Krause *et al*, 2011; Lee *et al*, 2005; Windle *et al*, 1997; Wsol *et al*, 2008).

The dorsal motor nucleus of vagus (10N) in the medulla also displayed high *Oxtr* expression, as has been shown previously in the rat (Dreifuss *et al*, 1988; Raggenbass *et al*, 1988; Raggenbass *et al*, 1987). The OXT-sensitive vagal neurons were mostly preganglionic motor neurons, projecting to the cervical, thoracic and abdominal visceral areas (Raggenbass *et al*., 1987). It has also been shown that the microinjection of an OXT antagonist into the 10N blocks the increase in gastric acid secretion and bradycardia induced by electrical stimulation of the PVH-th-is suggests a role for central OXT in autonomic efferent activity (Rogers & Hermann, 1986).

Despite higher plasma OXT levels in women than in men (Marazziti *et al*, 2019), prior, largely immunohistochemistry-based studies failed to identify a sex difference in *Oxt* expression in the brain. Similar numbers of OXT-positive immunoreactive (-IR) neurons were found in the PVH, SON, MPOA and bed nucleus of stria terminalis (BNST) of prairie, pine, meadow and montane voles (Wang *et al*, 1996), PVH and SON of naked mole rats (Rosen *et al*, 2008), and PVH, MPOA, LH and anterior hypothalamus (AH) of long-tailed hamsters (Xu *et al*, 2010). Furthermore, no sex differences were detected in OXT-IR neurons in the PVH, SON, BNST, MeA in several species of non-human primates (Caffe *et al*, 1989; Wang *et al*, 1997a; Wang *et al*, 1997b). There were also no sex differences in *Oxt* mRNA expression in the PVH and SON of the rat [for review, see: (Dumais & Veenema, 2016)]. Lastly, there were no sex differences in the number or size of OXT neurons in the PVH and SON in humans (Fliers *et al*, 1985; Ishunina & Swaab, 1999; Wierda *et al*, 1991).

By contrast, here we establish sex differences in *Oxt* expression in the mouse brain. Both the hypothalamus and forebrain of the females contained approximately twice as many *Oxt-*positive cells compared with males. Whereas, as expected, hypothalamic PVH of both sexes had high *Oxt* expression, the medial preoptic area (MPA) of the female forebrain and the lateral preoptic area (LPO) of the male forebrain contained the highest number of *Oxt-*expressing cells.

Finally, we have recently published an atlas of pituitary glycoprotein hormone receptors, namely *Tshr*, *Fshr* and *Lhcgr*, in more than 400 brain sites (Ryu *et al*, 2022). Surprisingly, we find a striking overlap in receptor distribution between the four receptors, including the *Oxtr-*with highest transcript levels in the ependymal layer of the third ventricle and olfactory bulb. While the role of olfactory OXTRs in social recognition is well established (Oettl & Kelsch, 2018; Oettl *et al*, 2016; Sun *et al*, 2021), the functional significance of OXTRs in the ependymal layer is yet unknown. However, in light of ubiquitous and newly emerging OXTR expression in the brain and peripheral organs, ependymal OXTRs seem to have an important role in gating bidirectional brain-periphery crosstalk.

In all, studies on central OXT signaling and its control of reproductive, metabolic and ingestive functions, and social behaviors occupy the vast majority of the literature. It is our hope that this comprehensive compendium of *Oxtr* and gender-specific *Oxt* expression in the brain will stimulate further investigations by others. In more general terms, the direct mapping of receptor expression in the brain and periphery provides the framework for determining new functions of ancient pituitary hormones and helps refocus at least some in the field towards paradigm-shifting discoveries of non-traditional, multifaceted roles of OXT.

## METHODS

### Mice

Adult mice (∼3 to 4-month-old) were housed in a 12 h:12 h light:dark cycle at 22 ± 2 °C with *ad libitum* access to water and regular chow. All procedures were approved by the Mount Sinai Institutional Animal Care and Use Committee and are in accordance with Public Health Service and United States Department of Agriculture guidelines.

### RNAscope

Brains and pituitary glands were collected from male and female mice for RNAscope. Briefly, mice were anesthetized with isoflurane (2 to 3 % in oxygen; Baxter Healthcare, Deerfield, IL) and transcardiacally perfused with 0.9% heparinized saline followed by 4% paraformaldehyde (PFA). Brains and pituitary and adrenal glands were extracted, sectioned into 0.5-cm-thick slices and post-fixed in 4 % PFA for 12 hours before being transferred to 30% sucrose (w/v) for 48 hours at 4 °C and then frozen. Brain coronal sections were cut at 10 μ m (pituitary and adrenal glands -at 5 μm), with every tenth section mounted onto ∼20 slides with six sections on each slide. This method ensures the accurate interrogation of the entire tissue without counting the same transcript twice. Sections were stored at -80 °C until required.

Detection of mouse *Oxt* and *Oxtr* was performed separately on fixed, frozen sections using Advanced Cell Diagnostics (ACD) RNAscope 2.5 LS Multiplex Reagent Kit and two RNAscope 2.5 LS probes, namely Mm-OXT (#493178) and Mm-OXTR (#412178). Epididymis and liver served as positive and negative controls for *Oxtr*, respectively. For *Oxt*, magnocellular cells of the PVH and SON served as positive controls while the brain from *Oxt^-^*^/-^k nockout mouse served a negative control.

Slides were thawed at room temperature for 10 minutes before baking at 60 °C for 45 min. Sections were then post-fixed in pre-chilled 4 % PFA for 15 minutes at 4 °C, washed in 3 changes of PBS for 5 min each before dehydration through 50 %, 70 and 100 % ethanol for 5 minutes each. The slides were air-dried for 5 minutes before loading onto a Bond Rx instrument (Leica Biosystems). Slides were prepared using the frozen slide delay prior to pre-treatments using Epitope Retrieval Solution 2 (Leica Biosystems) at 95 °C for 5 minutes, and ACD Enzyme from the Multiplex Reagent kit at 40 °C for 10 minutes.

Probe hybridization and signal amplification were performed per manufacturer’s instructions for chromogenic assays. Slides were imaged on a CellDiscoverer 7 microscope (Zeiss). Z-stack images with 1.0 μm spacin representative slice from each mouse corresponding to 4.28 to -7.92 mm A/P from Bregma using a 25x water immersion objective. Z-stacks were deconvoluted and compressed into 2D images using extended depth of focus (EDF) with maximum projection processing (ZEN Blue, Zeiss). EDF images were read into HALO v2.3 (Indica Labs) as .CZI files for analysis. OXT-and *Oxtr-*positive cells were detected using the CaseViewer 2.4 (3DHISTECH, Budapest, Hungary) or QuPath-0.2.3 (University of Edinburgh, UK) software based on OXT and its receptor intensity thresholds, size and shape.

## DATA AVAILABILITY

All data generated or analyzed during this study are included in the manuscript and supporting files; source data files have been provided for Figures 1, 2, 3 and Supplementary Figure 1.

## ACKNOWLEDGEMENTS

Work at Icahn School of Medicine at Mount Sinai carried at the Center for Translational Medicine and Pharmacology was supported by R01 AG071870, R01 AG074092 and U01 AG073148 to T.Y. and M.Z.; and U19 AG060917 and R01 DK113627 to M.Z. J.J.C. contributed to the concept and discussion of the study and was supported by the Agricultural Research Service of the United States Department of Agriculture, #3062-51000-053-00D. Mention of trade names or commercial products in this publication is solely for the purpose of providing specific information and does not imply recommendation or endorsement by the U.S. Department of Agriculture. The findings and conclusions in this manuscript are those of the authors and should not be construed to represent any official USDA of U.S. Government determination or policy.

## FIGURE LEGENDS

**Supplementary Figure 1:**
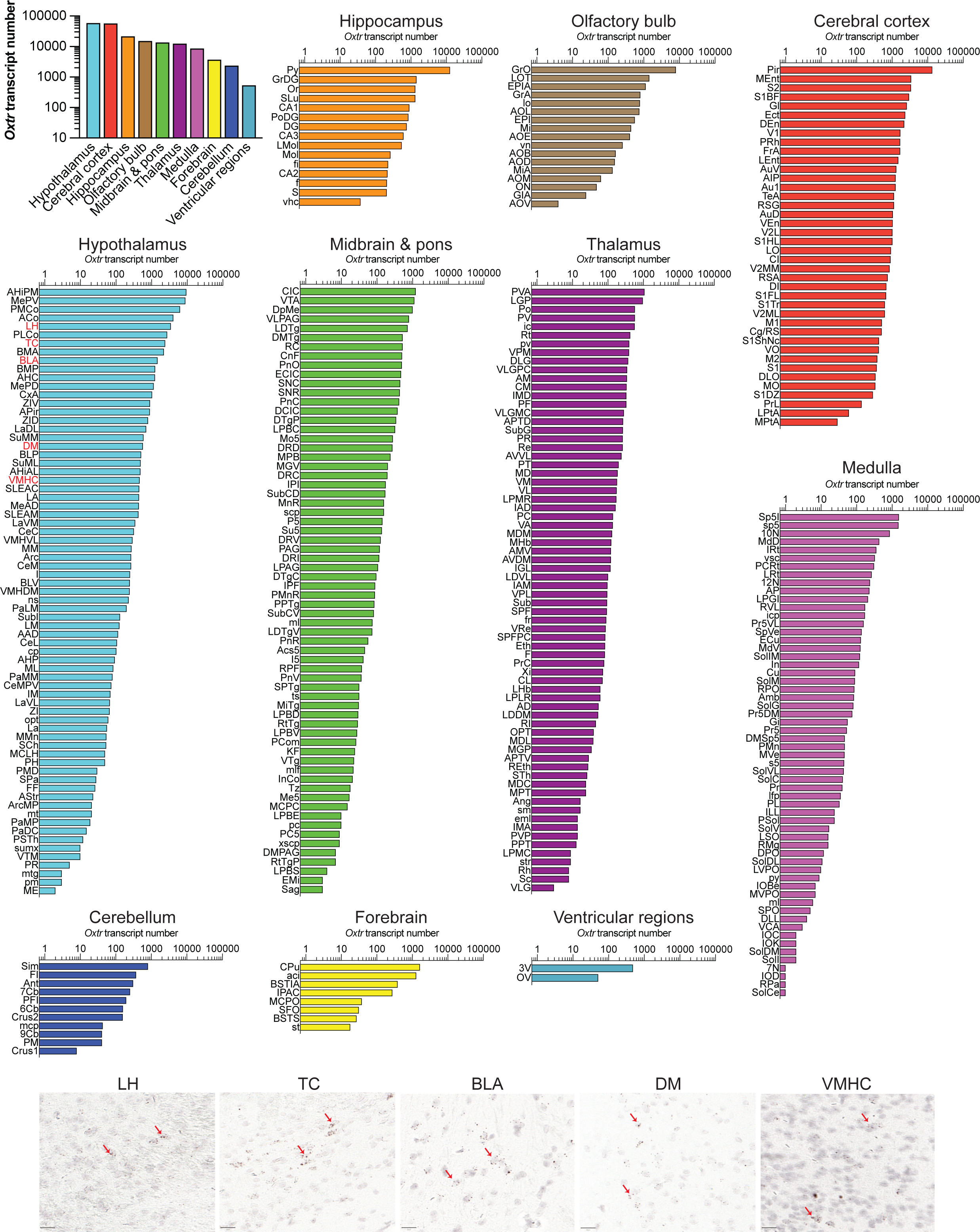
*Oxtr* expression in the mouse brain. *Oxtr* transcript number in brain regions and in nuclei, sub-nuclei and regions of the hypothalamus, cerebral cortex, hippocampus, olfactory bulb, midbrain and pons, thalamus, medulla, forebrain, cerebellum and ventricular system detected by RNAscope. Also shown are representative RNAscope images in several hypothalamic regions, including lateral hypothalamus (LH), tuber cinereum (TC), anterior part of the basolateral amygdaloid nucleus (BLA), dorsomedial hypothalamus (DM) and central part of ventromedial hypothalamic nucleus (VMHC). Note that LH and DM send SNS outflow to both bone and fat tissue. Scale bar: 20 µm.

## Appendix: Glossary of the brain nuclei, sub-nuclei and regions.

## REFERENCES

Abe E, Marians RC, Yu W, Wu XB, Ando T, Li Y, Iqbal J, Eldeiry L, Rajendren G, Blair HC, Davies TF, Zaidi M (2003) TSH is a negative regulator of skeletal remodeling. Cell 115: 151–162

Adan RA, Van Leeuwen FW, Sonnemans MA, Brouns M, Hoffman G, Verbalis JG, Burbach JP (1995) Rat oxytocin receptor in brain, pituitary, mammary gland, and uterus: partial sequence and immunocytochemical localization. Endocrinology 136: 4022–4028

Bale TL, Davis AM, Auger AP, Dorsa DM, McCarthy MM (2001) CNS region-specific oxytocin receptor expression: importance in regulation of anxiety and sex behavior. J Neurosci 21: 2546–2552

Bale TL, Dorsa DM (1995a) Regulation of oxytocin receptor messenger ribonucleic acid in the ventromedial hypothalamus by testosterone and its metabolites. Endocrinology 136: 5135–5138

Bale TL, Dorsa DM (1995b) Sex differences in and effects of estrogen on oxytocin receptor messenger ribonucleic acid expression in the ventromedial hypothalamus. Endocrinology 136: 27–32

Bohus B, Koolhaas JM, Korte SM, Roozendaal B, Wiersma A (1996) Forebrain pathways and their behavioural interactions with neuroendocrine and cardiovascular function in the rat. Clin Exp Pharmacol Physiol 23: 177–182

Bourque CW, Oliet SH, Kirkpatrick K, Richard D, Fisher TE (1993) Extrinsic and intrinsic modulatory mechanisms involved in regulating the electrical activity of supraoptic neurons. Ann N Y Acad Sci 689: 512–519

Caffe AR, Van Ryen PC, Van der Woude TP, Van Leeuwen FW (1989) Vasopressin and oxytocin systems in the brain and upper spinal cord of Macaca fascicularis. J Comp Neurol 287: 302–325

Caldwell JD, Prange AJ, Jr., Pedersen CA (1986) Oxytocin facilitates the sexual receptivity of estrogen-treated female rats. Neuropeptides 7: 175–189

Dreifuss JJ, Raggenbass M, Charpak S, Dubois-Dauphin M, Tribollet E (1988) A role of central oxytocin in autonomic functions: its action in the motor nucleus of the vagus nerve. Brain Res Bull 20: 765–770

Dubois-Dauphin M, Pevet P, Barberis C, Tribollet E, Dreifuss JJ (1992) Localization of binding sites for oxytocin in the brain of the golden hamster. Neuroreport 3: 797–800

Dumais KM, Veenema AH (2016) Vasopressin and oxytocin receptor systems in the brain: Sex differences and sex-specific regulation of social behavior. Front Neuroendocrinol 40: 1–23

Dyball RE, Paterson AT (1983) Neurohypophysial hormones and brain function: the neurophysiological effects of oxytocin and vasopressin. Pharmacol Ther 20: 419–436

Elands J, Beetsma A, Barberis C, de Kloet ER (1988) Topography of the oxytocin receptor system in rat brain: an autoradiographical study with a selective radioiodinated oxytocin antagonist. J Chem Neuroanat 1: 293–302

Fahrbach SE, Morrell JI, Pfaff DW (1984) Oxytocin induction of short-latency maternal behavior in nulliparous, estrogen-primed female rats. Horm Behav 18: 267–286

Fenselau H, Campbell JN, Verstegen AM, Madara JC, Xu J, Shah BP, Resch JM, Yang Z, Mandelblat-Cerf Y, Livneh Y, Lowell BB (2017) A rapidly acting glutamatergic ARC-->PVH satiety circuit postsynaptically regulated by alpha-MSH. Nat Neurosci 20: 42–51

Ferguson JN, Aldag JM, Insel TR, Young LJ (2001) Oxytocin in the medial amygdala is essential for social recognition in the mouse. J Neurosci 21: 8278–8285

Fliers E, Swaab DF, Pool CW, Verwer RW (1985) The vasopressin and oxytocin neurons in the human supraoptic and paraventricular nucleus; changes with aging and in senile dementia. Brain Res 342: 45–53

Grippo AJ, Trahanas DM, Zimmerman RR, 2nd, Porges SW, Carter CS (2009) Oxytocin protects against negative behavioral and autonomic consequences of long-term social isolation. Psychoneuroendocrinology 34: 1542–1553

Inada K, Tsujimoto K, Yoshida M, Nishimori K, Miyamichi K (2022) Oxytocin signaling in the posterior hypothalamus prevents hyperphagic obesity in mice. Elife 11

Insel TR (1990) Regional changes in brain oxytocin receptors post-partum: time-course and relationship to maternal behaviour. J Neuroendocrinol 2: 539–545

Insel TR (1992) Oxytocin--a neuropeptide for affiliation: evidence from behavioral, receptor autoradiographic, and comparative studies. Psychoneuroendocrinology 17: 3–35

Insel TR, Young L, Wang Z (1997) Central oxytocin and reproductive behaviours. Rev Reprod 2: 28–37

Ishunina TA, Swaab DF (1999) Vasopressin and oxytocin neurons of the human supraoptic and paraventricular nucleus: size changes in relation to age and sex. J Clin Endocrinol Metab 84: 4637–4644

Japundzic-Zigon N (2013) Vasopressin and oxytocin in control of the cardiovascular system. Curr Neuropharmacol 11: 218–230

Kombian SB, Mouginot D, Pittman QJ (1997) Dendritically released peptides act as retrograde modulators of afferent excitation in the supraoptic nucleus in vitro. Neuron 19: 903–912

Krause EG, de Kloet AD, Flak JN, Smeltzer MD, Solomon MB, Evanson NK, Woods SC, Sakai RR, Herman JP (2011) Hydration state controls stress responsiveness and social behavior. J Neurosci 31: 5470–5476

Landgraf R, Neumann ID (2004) Vasopressin and oxytocin release within the brain: a dynamic concept of multiple and variable modes of neuropeptide communication. Front Neuroendocrinol 25: 150–176

Lee PR, Brady DL, Shapiro RA, Dorsa DM, Koenig JI (2005) Social interaction deficits caused by chronic phencyclidine administration are reversed by oxytocin. Neuropsychopharmacology 30: 1883–1894

Ludwig M, Leng G (2006) Dendritic peptide release and peptide-dependent behaviours. Nat Rev Neurosci 7: 126–136

Marazziti D, Baroni S, Mucci F, Piccinni A, Moroni I, Giannaccini G, Carmassi C, Massimetti E, Dell’Osso L (2019) Sex-Related Differences in Plasma Oxytocin Levels in Humans. Clin Pract Epidemiol Ment Health 15: 58–63

Mitre M, Marlin BJ, Schiavo JK, Morina E, Norden SE, Hackett TA, Aoki CJ, Chao MV, Froemke RC (2016) A Distributed Network for Social Cognition Enriched for Oxytocin Receptors. J Neurosci 36: 2517–2535

Moos F, Richard P (1989) Paraventricular and supraoptic bursting oxytocin cells in rat are locally regulated by oxytocin and functionally related. J Physiol 408: 1–18

Nakahara TS, Camargo AP, Magalhaes PHM, Souza MAA, Ribeiro PG, Martins-Netto PH, Carvalho VMA, Jose J, Papes F (2020) Peripheral oxytocin injection modulates vomeronasal sensory activity and reduces pup-directed aggression in male mice. Sci Rep 10: 19943

Neumann I, Douglas AJ, Pittman QJ, Russell JA, Landgraf R (1996) Oxytocin released within the supraoptic nucleus of the rat brain by positive feedback action is involved in parturition-related events. J Neuroendocrinol 8: 227–233

Neumann I, Koehler E, Landgraf R, Summy-Long J (1994) An oxytocin receptor antagonist infused into the supraoptic nucleus attenuates intranuclear and peripheral release of oxytocin during suckling in conscious rats. Endocrinology 134: 141–148

Neumann I, Russell JA, Landgraf R (1993) Oxytocin and vasopressin release within the supraoptic and paraventricular nuclei of pregnant, parturient and lactating rats: a microdialysis study. Neuroscience 53: 65–75

Neumann ID (2007) Stimuli and consequences of dendritic release of oxytocin within the brain. Biochem Soc Trans 35: 1252–1257

Oettl LL, Kelsch W (2018) Oxytocin and Olfaction. Curr Top Behav Neurosci 35: 55–75

Oettl LL, Ravi N, Schneider M, Scheller MF, Schneider P, Mitre M, da Silva Gouveia M, Froemke RC, Chao MV, Young WS, Meyer-Lindenberg A, Grinevich V, Shusterman R, Kelsch W (2016) Oxytocin Enhances Social Recognition by Modulating Cortical Control of Early Olfactory Processing. Neuron 90: 609–621

Paxinos G, Franklin KBJ (2007) The Mouse Brain in Stereotaxic Coordinates. Academic Press, New York

Pedersen CA, Ascher JA, Monroe YL, Prange AJ, Jr. (1982) Oxytocin induces maternal behavior in virgin female rats. Science 216: 648–650

Pow DV, Morris JF (1989) Dendrites of hypothalamic magnocellular neurons release neurohypophysial peptides by exocytosis. Neuroscience 32: 435–439

Raggenbass M, Charpak S, Dubois-Dauphin M, Dreifuss JJ (1988) Electrophysiological evidence for oxytocin receptors on neurones located in the dorsal motor nucleus of the vagus nerve in the rat brainstem. J Recept Res 8: 273–282

Raggenbass M, Dubois-Dauphin M, Charpak S, Dreifuss JJ (1987) Neurons in the dorsal motor nucleus of the vagus nerve are excited by oxytocin in the rat but not in the guinea pig. Proc Natl Acad Sci U S A 84: 3926–3930

Rogers RC, Hermann GE (1986) Hypothalamic paraventricular nucleus stimulation-induced gastric acid secretion and bradycardia suppressed by oxytocin antagonist. Peptides 7: 695–700

Rosen GJ, de Vries GJ, Goldman SL, Goldman BD, Forger NG (2008) Distribution of oxytocin in the brain of a eusocial rodent. Neuroscience 155: 809–817

Ryu V, Garretson JT, Liu Y, Vaughan CH, Bartness TJ (2015) Brown adipose tissue has sympathetic-sensory feedback circuits. J Neurosci 35: 2181–2190

Ryu V, Gumerova A, Korkmaz F, Kang SS, Katsel P, Miyashita S, Kannangara H, Cullen L, Chan P, Kuo T, Padilla A, Sultana F, Wizman SA, Kramskiy N, Zaidi S, Kim SM, New MI, Rosen CJ, Goosens KA, Frolinger T et al (2022) Brain atlas for glycoprotein hormone receptors at single-transcript level. Elife 11

Ryu V, Watts AG, Xue B, Bartness TJ (2017) Bidirectional crosstalk between the sensory and sympathetic motor systems innervating brown and white adipose tissue in male Siberian hamsters. Am J Physiol Regul Integr Comp Physiol 312: R324–R337

Sato K, Hamasaki Y, Fukui K, Ito K, Miyamichi K, Minami M, Amano T (2020) Amygdalohippocampal Area Neurons That Project to the Preoptic Area Mediate Infant-Directed Attack in Male Mice. J Neurosci 40: 3981–3994

Shibuya I, Kabashima N, Ibrahim N, Setiadji SV, Ueta Y, Yamashita H (2000) Pre- and postsynaptic modulation of the electrical activity of rat supraoptic neurones. Exp Physiol 85 Spec No: 145S–151S

Sofroniew MV (1983) Morphology of vasopressin and oxytocin neurones and their central and vascular projections. Prog Brain Res 60: 101–114

Sun C, Yin Z, Li BZ, Du H, Tang K, Liu P, Hang Pun S, Lei TC, Li A (2021) Oxytocin modulates neural processing of mitral/tufted cells in the olfactory bulb. Acta Physiol (Oxf) 231: e13626

Sun L, Lizneva D, Ji Y, Colaianni G, Hadelia E, Gumerova A, Ievleva K, Kuo TC, Korkmaz F, Ryu V, Rahimova A, Gera S, Taneja C, Khan A, Ahmad N, Tamma R, Bian Z, Zallone A, Kim SM, New MI et al (2019) Oxytocin regulates body composition. Proc Natl Acad Sci U S A

Sun L, Tamma R, Yuen T, Colaianni G, Ji Y, Cuscito C, Bailey J, Dhawan S, Lu P, Calvano CD, Zhu LL, Zambonin CG, Di Benedetto A, Stachnik A, Liu P, Grano M, Colucci S, Davies TF, New MI, Zallone A, Zaidi M (2016) Functions of vasopressin and oxytocin in bone mass regulation. Proc Natl Acad Sci U S A 113: 164–169

Swanson LW, Sawchenko PE (1983) Hypothalamic integration: organization of the paraventricular and supraoptic nuclei. Annu Rev Neurosci 6: 269–324

Tamma R, Colaianni G, Zhu LL, DiBenedetto A, Greco G, Montemurro G, Patano N, Strippoli M, Vergari R, Mancini L, Colucci S, Grano M, Faccio R, Liu X, Li J, Usmani S, Bachar M, Bab I, Nishimori K, Young LJ et al (2009) Oxytocin is an anabolic bone hormone. Proc Natl Acad Sci U S A 106: 7149–7154

Tamma R, Sun L, Cuscito C, Lu P, Corcelli M, Li J, Colaianni G, Moonga SS, Di Benedetto A, Grano M, Colucci S, Yuen T, New MI, Zallone A, Zaidi M (2013) Regulation of bone remodeling by vasopressin explains the bone loss in hyponatremia. Proc Natl Acad Sci U S A 110: 18644–18649

Tribollet E, Dubois-Dauphin M, Dreifuss JJ, Barberis C, Jard S (1992) Oxytocin receptors in the central nervous system. Distribution, development, and species differences. Ann N Y Acad Sci 652: 29–38

van Wimersma Greidanus TB, Maigret C (1996) The role of limbic vasopressin and oxytocin in social recognition. Brain Res 713: 153–159

Wang Z, Moody K, Newman JD, Insel TR (1997a) Vasopressin and oxytocin immunoreactive neurons and fibers in the forebrain of male and female common marmosets (Callithrix jacchus). Synapse 27: 14–25

Wang Z, Toloczko D, Young LJ, Moody K, Newman JD, Insel TR (1997b) Vasopressin in the forebrain of common marmosets (Callithrix jacchus): studies with in situ hybridization, immunocytochemistry and receptor autoradiography. Brain Res 768: 147–156

Wang Z, Zhou L, Hulihan TJ, Insel TR (1996) Immunoreactivity of central vasopressin and oxytocin pathways in microtine rodents: a quantitative comparative study. J Comp Neurol 366: 726–737

Wierda M, Goudsmit E, Van der Woude PF, Purba JS, Hofman MA, Bogte H, Swaab DF (1991) Oxytocin cell number in the human paraventricular nucleus remains constant with aging and in Alzheimer’s disease. Neurobiol Aging 12: 511–516

Windle RJ, Shanks N, Lightman SL, Ingram CD (1997) Central oxytocin administration reduces stress-induced corticosterone release and anxiety behavior in rats. Endocrinology 138: 2829–2834

Wsol A, Cudnoch-Jedrzejewska A, Szczepanska-Sadowska E, Kowalewski S, Puchalska L (2008) Oxytocin in the cardiovascular responses to stress. J Physiol Pharmacol 59 Suppl 8: 123–127

Xu L, Pan Y, Young KA, Wang Z, Zhang Z (2010) Oxytocin and vasopressin immunoreactive staining in the brains of Brandt’s voles (Lasiopodomys brandtii) and greater long-tailed hamsters (Tscherskia triton). Neuroscience 169: 1235–1247

Yao S, Bergan J, Lanjuin A, Dulac C (2017) Oxytocin signaling in the medial amygdala is required for sex discrimination of social cues. Elife 6

Yoshimura R, Kiyama H, Kimura T, Araki T, Maeno H, Tanizawa O, Tohyama M (1993) Localization of oxytocin receptor messenger ribonucleic acid in the rat brain. Endocrinology 133: 1239–1246

Zaidi M, New MI, Blair HC, Zallone A, Baliram R, Davies TF, Cardozo C, Iqbal J, Sun L, Rosen CJ, Yuen T (2018) Actions of pituitary hormones beyond traditional targets. J Endocrinol 237: R83–R98

